# IGF2BP3 promotes the progression of gastric cancer by activating cGMP-PKG signaling pathway via targeting FBXO32

**DOI:** 10.1101/2024.06.28.601102

**Authors:** Yi Si, Bo Tian, Rui Zhang, Mingda Xuan, Kunyi Liu, Jiao Jiao, Shuangshuang Han, Hongfei Li, Yanhong Hu, Hongyan Zhao, Wenjing He, Jia Wang, Ting Liu, Weifang Yu

**Author notes:** **Corresponding author:** Weifang Yu, M.D., Professor, No. 89 Donggang Road, Shijiazhuang, Hebei, China. Ting Liu, No. 89 Donggang Road, Shijiazhuang, Hebei, China. Jia Wang, Professor, No. 89 Donggang Road, Shijiazhuang, Hebei, China. These authors contributed equally (Yi Si, Bo Tian).

## Abstract

N6-methyladenosine (m^6^A) represents the most prevalent chemical modification on eukaryotic mRNA, with an accumulating body of literature indicating its pivotal significance in the pathogenesis of human cancers. Nevertheless, the precise molecular interplay between the m^6^A reader protein IGF2BP3 and gastric cancer remains to be thoroughly delineated. Our study uncovered that the expression of IGF2BP3 in gastric cancer tissues is markedly elevated in comparison to adjacent normal tissues, and this upregulation is tightly correlated with the incidence of lymph node metastasis, more advanced TNM stages, and deeper invasion depth of tumor in patients. *In vitro* experiments demonstrated that IGF2BP3 potentiates the proliferative, migratory, and invasive capacities of gastric cancer cells, while concurrently inhibiting apoptosis and augmenting the intracellular levels of aerobic glycolysis. *In vivo* experiments revealed that IGF2BP3 contributes to the growth of gastric cancer. Mechanistically, IGF2BP3 can increase the expression of FBXO32 protein by recognizing and binding to the m^6^A binding site on FBXO32 mRNA and further activate the downstream cGMP-PKG signaling pathway, thereby modulating various biological functions of gastric cancer cells and ultimately promoting the progression of gastric cancer. In summary, our findings suggest that IGF2BP3 upregulates the expression of FBXO32 protein in an m^6^A dependent manner and subsequently activates the cGMP-PKG signaling pathway, ultimately leading to the onset and progression of gastric cancer. Consequently, the targeting of the IGF2BP3/FBXO32/cGMP-PKG axis emerges as a promising therapeutic modality for the treatment of gastric cancer.

## Introduction

Gastric cancer (GC) is a heterogeneous disease characterized by significant differences in epidemiology and histopathology across countries and is one of the leading causes of cancer-related deaths^[1]^. The morbidity and mortality rates of GC are high worldwide and particularly in China, posing a serious threat to human health^[2]^. Typically, there are no obvious clinical symptoms in the early stage of GC, and most patients are diagnosed at an advanced stage. The recurrence rate of advanced GC is high, leading to a poor overall prognosis for GC patients^[3]^. To date, the specific molecular mechanisms of GC have not been fully elucidated. This may be attributed to its complex etiology, which primarily involves the interactions between oncogenes and tumor suppressor genes^[4]^, as well as genetic, epigenetic, and environmental factors^[5]^. Therefore, further exploration of the molecular mechanisms underlying the occurrence and development of GC will provide potential new targets for the clinical treatment of GC, which is of significant clinical importance.

In recent years, the role of epigenetics in the pathogenesis of GC has attracted extensive attention from scholars worldwide. Epigenetics is generally defined as the study of chemical modifications to DNA and RNA, which mainly include DNA and RNA methylation, histone modification, non-coding RNA modification, and chromatin rearrangement^[6]^. N6-methyladenosine (m^6^A) modification is an integral part of epigenetics research. m^6^A modification is the most common internal modification of eukaryotic mRNA, regulating the stability, translation efficiency, alternative splicing and localization of RNA at the post-transcriptional level. It participates in various biological processes such as initiation, proliferation, differentiation, metastasis and metabolic reprogramming of tumor cells^[7, 8]^. m^6^A related proteins are mainly divided into methyltransferase, demethylase and reader proteins, involving 21 related genes^[9]^. By analyzing the expression of m^6^A related proteins in GC and adjacent normal tissues in the TCGA and GEO (GSE54129 and GSE66229 data sets) databases, we screened out four ‘readers’, YTHDF1, YTHDF2, IGF2BP1 and IGF2BP3, whose expression levels were higher in cancer tissues than in adjacent normal tissues in the above databases. Among these, the expression difference of IGF2BP3 was the most significant, and there are few studies about IGF2BP3 and GC, so IGF2BP3 was selected as the research target. It is reported that the main function of m^6^A ‘readers’ is to interpret RNA methylation modification information and regulate the translation and degradation of its downstream RNA^[10, 11]^.

Insulin like growth factor II mRNA binding protein 3 (IGF2BP3), an m^6^A ‘reader’, is located on human chromosome 7p15.3 and contains two RNA recognition motifs and four K homologous domains^[12]^. Overexpression of IGF2BP3 was first observed in pancreatic cancer^[13]^. Its primary role is to recognize m^6^A binding sites through the K homology domain, thereby increasing the stability and translation efficiency of target mRNAs^[14]^. IGF2BP3 is implicated in the development of various cancers, playing distinct roles in each, such as, IGF2BP3 enhances the stability of TMBIM6 mRNA by identifying the m^6^A site, thus promoting the progression of laryngeal squamous cell carcinoma^[15]^; Similarly, IGF2BP3 stabilizes lnc-CTHCC by recognizing its m^6^A site, which increases lnc-CTHCC expression in hepatocellular carcinoma, ultimately facilitating the disease’s progression^[16]^; In colorectal cancer, IGF2BP3 stabilizes CCND1 mRNA by identifying its m^6^A site, promoting CCND1 protein expression, regulating the cell cycle, and thus promoting the proliferation and growth of cancer cells^[17]^; Furthermore, Wang et al. found that m^6^A ‘reader’ IGF2BP2 enhances the stability of GLUT1 mRNA, a key gene in aerobic glycolysis, in an m^6^A dependent manner, significantly promoting the aerobic glycolysis of colorectal cancer cells^[18]^. Therefore, IGF2BP3 may be a critical oncogene. We observed that IGF2BP3 was highly expressed in GC and was closely related to poor prognosis in GC patients. IGF2BP3 reinforces the malignant phenotype of GC cells and promotes the growth of GC *in vivo*. Through omics analysis and various experiments, we identified for the first time that the downstream gene with m^6^A modification of IGF2BP3 in GC is FBXO32, which is also highly expressed in GC and interacts with IGF2BP3. IGF2BP3 upregulates the expression of FBXO32 protein in an m^6^A dependent manner and activates the cGMP-PKG signaling pathway, which further enhances the proliferation, migration, and invasion capabilities of GC cells, and ultimately promotes the progression of GC.

## Materials and Methods

### Patients and samples

63 pairs of GC tissues and adjacent normal tissues were obtained from the First Hospital of Hebei Medical University. None of the patients had received radiation or chemotherapy prior to surgery and they had signed informed consent forms. The above tissue samples were promptly placed in liquid nitrogen after isolation, and then transferred to the Biological Sample Bank of the First Hospital of Hebei Medical University for storage. The cancer tissues used in this study were pathologically confirmed as gastric adenocarcinoma/signet-ring cell carcinoma. This study was approved by the Ethics Committee of the First Hospital of Hebei Medical University (Approval letter No. S00180) and adheres to the principles of the Declaration of Helsinki.

### Cell culture

Human gastric mucosa cell (GES-1) and GC cells (HGC-27, MKN7, MKN74) were obtained from Wuhan Pricella Biotechnology Co., Ltd., and were verified by STR analysis. The cells were cultured in RPMI 1640 medium (Gibco, Gaithersburg, MD, USA) supplemented with 10% fetal bovine serum (FBS; Gibco) and 1% penicillin-streptomycin solution (Solarbio Sciences & Technology Co. Ltd, Beijing, China). They were maintained in a constant temperature and humidity incubator with 5% carbon dioxide at 37℃, and routine mycoplasma detection was performed monthly using PCR. In all experiments, the cells were used for a maximum of no more than 25 passages.

### Plasmids, siRNAs, shRNAs, Transfections and Stable cell lines

Transfection was performed according to Lipofectamine 3000 (Invitrogen, Carlsbad, CA, USA) when the cell confluence reached 80-90% in a cell culture flask or dish. IGF2BP3 siRNA and negative control (GenePharma Co., Ltd., Shanghai, China) were transfected into HGC-27 and MKN74 cells, while an IGF2BP3 overexpression plasmid and a negative control (GenePharma) were transfected into MKN7 and GES-1 cells. Additionally, IGF2BP3 shRNA and a negative control (GenePharma) were transfected into HGC-27 cells. FBXO32 siRNA and a negative control (GenePharma) were transfected into MKN7 cells, and an FBXO32 overexpression plasmid and a negative control (GenePharma) were transfected into HGC-27 cells. Follow-up experiments were conducted 48 hours after transfection. To eatablish stable transfected cell lines (HGC-27 shNC, HGC-27 shIGF2BP3), the cell culture medium was changed to include neomycin at a final concentration of 600 μg/mL (without other antibiotics) 48 hours after transfection with IGF2BP3 shRNA in HGC-27 cells. Once the clones have formed, the neomycin concentration was adjusted to 300 μg/mL, continue culturing until day 10.

### RNA extraction and qRT-PCR

Total RNA was extracted from cells and tissues using RNA-Easy Isolation Reagent (Vazyme Biotech Co.,Ltd., Nanjing, China). The RNA was then reverse-transcribed into complementary DNA (cDNA) following the instructions of PrimeScript RT reagent kit (Takara, Beijing, China). qRT-PCR was performed using the AceQ Universal SYBR qPCR Master Mix (Vazyme), with β-actin serving as the endogenous control mRNA. Each sample was tested in triplicate. The mRNA expression level was determined based on the CT (cyclic threshold) value of each sample, and the relative expression level of each sample was calculated using the 2^-ΔΔCT method. The sequence of primers are detailed in Supplementary Table 1.

### Immunohistochemistry (IHC)

Tissues from patients and nude mice were fixed with 4% paraformaldehyde, followed by paraffin embedding (Shanghai YiYang Instrument Co., Ltd., Shanghai, China), sectioning, and staining. The Rabbit/Mouse two-step detection kit (Beijing ZSBG-Bio Co., Ltd., Beijing, China) was used for immunohistochemical analysis according to the manufacturer’s instructions. Anti-IGF2BP3 (diluted 1:150; Proteintech, Wuhan, Hubei, P.R.C; Catalog number: 14642-1-AP), Anti-Ki67 (diluted 1:100; Proteintech; Catalog number: 27309-1-AP) and Anti-FBXO32 (diluted 1:100; Proteintech; Catalog number: 67172-1-Ig) antibodies were used for immunohistochemical staining. The staining results were analyzed by using a double-blind scoring method and the average optical density (AOD) method.

### Western blot analysis and antibodies

Protein was extracted using RIPA lysis buffer (Solarbio) and the protease/phosphatase inhibitor (Solarbio) at a ratio of 100:1. The proteins were separated electrophoretically on a 10% SDS polyacrylamide gel (Bio-Rad Laboratories Inc, Hercules, CA, USA) and transferred to a polyvinylidene fluoride membrane (Merck Millipore, Billerica, MA, USA). The following antibodies were used in this study: IGF2BP3 antibody (diluted 1:1000; Abcam, Cambridge, MA, USA; Catalog number: ab177477), β-actin antibody (diluted 1:1500; ZSBG-Bio; Catalog number: TA-09), FBXO32 antibody (diluted 1:500; Abcam; Catalog number: ab168372) and GAPDH antibody (diluted 1:500; Goodhere Co. Ltd, Hangzhou, China; Catalog number: AB-P-R 001), PRKG1 (PKG1) Polyclonal antibody (diluted 1:100; Proteintech; Catalog number: 21646-1-AP), Phospho-VASP (Ser239) Antibody (diluted 1:100; Affinity; Catalog number: AF3338), VASP Polyclonal antibody (diluted 1:100; Proteintech; Catalog number: 13472-1-AP). The immunoreactive protein bands were detected by the Odyssey Scanning System (LI-COR Biosciences, Lincoln, NE, USA).

### Cell proliferation assay

For CCK-8 assay, the cells of each group were uniformly distributed in 96-well plates with a density of 1×10^3^ cells per well, and cultured with complete culture medium. At 24, 48, 72, and 96 hours after distribution, 10 μL of Cell Counting Kit −8 (CCK-8; Dojindo, Tokyo, Japan) reagent was added to each well. Then, the cells were incubated in the incubator for an additional 2 hours. The absorbance of each well at 450 nm was measured using a Promega GloMax luminescence detector (Promega, Madison, WI, USA).

For the colony formation assay, cells of each group were uniformly distributed in 6-well plates at a density of 500 cells per well. Fresh culture medium was replaced every 3 days. Cells were cultured for 7-14 days, then washed twice with PBS, treated with 4% paraformaldehyde for 30 minutes, and stained with 0.1% crystal violet for 20 minutes.

### Cell migration assay

For the wound healing assay, the cells of each group were uniformly distributed in 6-well plates. When the cell confluence reached or was close to 100%, a 200 μL pipette tip was used to draw two straight lines on each well to simulate the wound. The cells were then washed twice with PBS and cultured with a culture medium without FBS. Each well was photographed at 0 hours and 48 hours after the wound was created, and the migration rate was expressed as the ratio of gap width measured at 48 hours to the gap width measured at 0 hour.

For the Transwell migration assay, 3×10^4^ (HGC-27, GES-1) /1×10^5^ (MKN7, MKN74) cells were added to the upper chamber (Corning Incorporated, Corning, NY, USA) in 200 μL FBS-free medium. To induce downward migration, 700 μL of complete medium was added to the lower chamber. After 48 hours of culture, cells on the upper chamber side of the polycarbonate membrane were removed using a swab. The chamber was cleaned twice with PBS, then treated with 4% paraformaldehyde for 30 minutes, followed by 0.1% crystal violet for 20 minutes. The excess crystal violet was cleaned again with PBS. Five visual fields were randomly selected for photography, and the stained cells were counted using ImageJ software (National Institutes of Health, Bethesda, MD, USA).

### Cell invasion assay

For the Transwell invasion assay, cells were resuspended in 100 μL FBS-free medium and added to the upper chamber, while 500 μL of complete medium was added to the lower chamber to induce downward invasion of the cells in the upper chamber. All other experimental procedures were the same as those for the Transwell migration assay.

### Cell apoptosis assay

Cells from each group were collected 48 hours after transfection and stained using the Annexin V-FITC/PI apoptosis assay kit (NeoBioscience, Shenzhen, China). Flow cytometry (BD Biosciences, San Jose, CA, USA) was used to analyze the apoptosis of 1×10^5^ cells, and FlowJo software was used to calculate the percentage of apoptotic cells.

### Cell glucose metabolism assay

MKN7/HGC-27 cells that overexpressed or have knockdown of IGF2BP3 were collected and processed according to the kit instructions (Glucose Assay kit, Abcam, ab65333; L-Lactate Assay kit, Abcam, ab65331; ATP Assay Kit, Abcam, ab83355). In brief, the standard solutions of glucose, lactic acid and ATP were prepared in a gradient dilution. The reaction mixtures were then prepared, and the diluted standard solutions were mixed with the reaction mixtures for incubation (37℃ or room temperature) for 30 minutes. The spectrophotometer was adjusted to the corresponding wavelength to read the absorbance value, and the standard curve for glucose, lactic acid and ATP were generated. The corresponding reagents from the kits were added to the lysed cells, and the cell lysate was deproteinized using Deproteinizing Sample Preparation Kit - TCA (Abcam; Catalog number: ab204708). The deproteinized cell lysates were then mixed with the reaction mixtures for incubation (37℃ or room temperature) for 30 minutes, and the absorbance values were compated to the standard curves to calculate the concentrations of glucose, lactic acid and ATP.

### Animal studies

Ten male BALB/c nude mice aged 4-6 weeks, 16-20 g (procured from Beijing Huafukang Biotechnology Co., LTD.) were randomly divided into two groups (five mice per group) and housed in a pathogen-free environment with sufficient water and food. The room temperature was maintained at approximately 22℃ and a 12/12-hour light/dark cycle. In a clean bench, the skin of the left hind limb flank of the nude mice was disinfected, and HGC-27 cells (5×10^6^ cells per mice) stably transfected with shRNA-NC and shRNA-IGF2BP3 were subcutaneously injected using a sterile syringe. Starting from day 3 post-injection, tumor size was evaluated every two days using the formula: volume = (long diameter × short diameter ^2^)/2. When the tumor volume approached 1000 mm^3^, the nude mice were euthanized humanely. Tumor specimens were collected, photographed, weighed, and fixed in formalin. The tumor samples were then paraffin-embedded and sectioned for subsequent HE and IHC analysis. The above experiments were approved by the Ethics Committee of the First Hospital of Hebei Medical University and carried out in accordance with the experimental animal care and use system.

### RNA sequencing (RNA-seq)

Total RNA was extracted from HGC-27 cells transfected with siRNA-NC and siRNA-IGF2BP3 for 24 hours. The RNA library construction and sequencing analysis were entrusted to Beijing Novogene Technology Co., LTD. In short, a total of six samples (siNC× 3, siIGF2BP3 × 3) were tested using Illumina’s NovaSeq 6000 platform. Quality control and filtering are carried out on the output data from each sample to obtain high-quality data. HISAT software was used to align high-quality data to reference genomes and to quantify gene or transcript expression levels in the sample. DESeq2 software was used for differential expression analysis, with differential genes defined as those with log_2_|Fold Change| > 1.0 and *P* < 0.05 were selected.

### Methylated RNA immunoprecipitation sequencing (MeRIP-seq)

Total RNA was extracted from HGC-27 cells transfected with siRNA-NC and siRNA-IGF2BP3 for 24 hours. The RNA library construction and sequencing analysis were entrusted to Novogene. In short, the original data was processed using fastp (version 0.19.11) to obtain high quality data. The reference genome index was constructed using BWA (version 0.7.12), and the high-quality data was aligned with the reference genome using BWA mem (version 0.7.12). After mapping reads to the reference genome, the m^6^A peaks within each group were identified using the exomePeak R package (version 2.16.0). The genes corresponding to each peak were presumed to be peak-related genes. The m^6^A enrichment motif of each group was identified by HOMER (version 4.9.1). The peak distribution in the 5’ UTR, CDS, 3’ UTR and other functional regions on mRNA transcripts was statistically analyzed.

### RNA immunoprecipitation and high-throughput sequencing assay (RIP-seq)

According to the instructions of the RNA Immunoprecipitation (RIP) Kit (BersinBio, Guangzhou, China; Catalog number: Bes5101), HGC-27 cells transfected with siRNA-NC and siRNA-IGF2BP3 for 48 hours were collected and lysed. An IGF2BP3 specific antibody (10 μg; Proteintech; Catalog number: 14642-1-AP) was employed to bind the endogenous expression of IGF2BP3 protein in the cells, precipitate the target protein-RNA complex, and subsequently isolate and purify the RNA within the complex. Novogene was commissioned to perform ribosomal RNA removal, RNA library construction and high-throughput sequencing analysis. In brief, RNA fragment distribution and concentration after immunoprecipitation and ribosomal RNA removal were measured using an Agilent 2100 bioanalyzer (Agilent) and a simpliNano spectrophotometer (GE Healthcare). The library was constructed using the NEB Next®Ultra™RNA Library Preparation Kit (New England Biolabs). The library quality was assessed using the Agilent Bioanalyzer 2100 system and then sequenced using the Illumina Novaseq platform. The fastp software was utilized to filter the original data and obtain high-quality data. The reference genome index was established using BWA (version 0.7.12), and the filtered data was compared with the reference genome using BWA mem (version 0.7.12). After the comparison was completed, the IP enrichment region was identified against the background using MACS2(version 2.1.0) peak call software. After peak calling was completed, the results were analyzed for chromosome distribution, peak width, fold abundance, significance level and peak number per peak. Homer (version 4.9.1) software was used to identify m^6^A enrichment motifs in each group. Peak Annotator software was used to identify peak-related genes. Differential peak analysis was carried out based on the different enrichment times of the peaks in the results, and then the genes related to the difference peak were identified.

### Co-Immunofluorescent (Co-IF) assay

HGC-27 cells were evenly distributed on coverslips in a 6-well plate. When the cell fusion rate reached 20-30%, the cells were gently rinsed with PBS for three times. Subsequently, the cells were fixed with 4% paraformaldehyde, permeated with 0.2% Triton X-10 (Solarbio; Catalog number: T8200), and blocked with 2% bovine serum albumin (BSA-V; Solarbio; Catalog number: A8020). The cells were then incubated overnight at 4℃ with IGF2BP3 antibody (diluted 1:50; Proteintech; Catalog number: 14642-1-AP) and FBXO32 antibody (diluted 1:50; Proteintech; Catalog number: 67172-1-Ig). Following this, the cells were incubated with fluorescent secondary antibody (diluted 1:500; Cy3 anti-rabbit IgG, FITC anti-mouse IgG; Beyotime Biotechnology Co., Shanghai, China) at room temperature for 1 hour in a dark environment. Finally, 4 ‘6’ -diamino-2-phenylindole dihydrochloride (DAPI; Beyotime Biotechnology Co.) was used to stain the nuclei. Representative images were captured using a laser scanning confocal microscope (ZEISS, Oberkochen, Germany).

### Bioinformatics analysis

The expression of IGF2BP3 mRNA in cancer and adjacent tissues of patients with GC was obtained from TCGA (https://www.aclbi.com/static/index.html#/tcga) and GEO (https://www.aclbi.com/static/index.html#/geo) databases. The online website Kaplan-Meier Plotter (https://kmplot.com/analysis/) was used to analyze the influence of gene expression level on the prognosis of GC patients. GEPIA2 (http://gepia2.cancer-pku.cn/#index) was used to analyze gene expression levels in GC and adjacent normal tissues, while GeneCards (https://www.genecards.org/) provided insight into gene sublocalization within cells. PrimerBank (https://pga.mgh.harvard.edu/primerbank/) and the Primer designing tool (https://www.ncbi.nlm.nih.gov/tools/primer-blast/) were employed for primer prediction. BioRender (https://www.biorender.com/) facilitated the creation of scientific schematic diagrams, and catRAPID omics (http://s.tartaglialab.com/page/catrapid_group) was used to predict binding site motifs between the IGF2BP3 protein and FBXO32 mRNA. The iGV software was utilized for the analysis of genomic m^6^A methylation levels.

### Statistical analysis

In this study, three independent replicates were performed for each group of experiments. The results of normal distribution data were presented as mean ± standard deviation, while non-normally distributed data were presented as median and interquartile distance. Statistical analysis in this study was processed using GraphPad Prism 9.5 (GraphPad Software, La Jolla, CA, USA) and SPSS 26.0 (IBM, Armonk, NY, USA). Statistical methods including Student’s *t*-test, one-way ANOVA and two-way ANOVA were employed. A significance level of *P*<0.05 was considered statistically significant.

### Data Availability

The data generated in this study are available upon request from the corresponding author. The data analyzed in this study were obtained from The Cancer Genome Atlas (TCGA) and Gene Expression Omnibus (GEO) at GSE54129 and GSE66229.

## Results

### IGF2BP3 is upregulated in GC patients and is associated with poor prognosis

To investigate the role of m^6^A related genes in the pathogenesis of GC, we first analyzed the expression of 21 m^6^A related genes in GC and adjacent normal tissues in the TCGA database and the GSE54129 and GSE66229 datasets (Figure 1A-C). Four m^6^A reader proteins (YTHDF1, YTHDF3, IGF2BP1 and IGF2BP3) with higher expression levels in GC than in adjacent normal tissues were screened. Among them, IGF2BP3 exhibited the most significant differential expression, and due to the limited research on its mechanism in GC, we selected IGF2BP3 as our research focus. Kaplan-Meier survival analysis suggested that GC patients with high IGF2BP3 expression had worse overall survival compared to those with low IGF2BP3 expression (Figure 1D). We then assessed IGF2BP3 mRNA expression in 50 pairs of fresh frozen GC tissues and adjacent normal tissues using qRT-PCR. Consistent with the TCGA and GEO data, the expression of IGF2BP3 was significantly higher in GC tissues compared to adjacent normal tissues, with a statistically significant difference (*P*<0.01) (Figure 1E). IHC was employed to detect IGF2BP3 protein expression in these samples. The results showed that IGF2BP3 protein was primarily localized in the cytoplasm of GC cells, and was expressed at significantly higher levels than in adjacent normal tissues (χ^2^ = 22.415), with a statistically significant difference (*P*<0.001) (Figure 1F, Table 1). Finally, we analyzed the correlation between IGF2BP3 expression and the clinical characteristics of GC patients. The results suggest that IGF2BP3 expression is significantly associated with lymph node metastasis (*P*<0.001), TNM staging (*P* = 0.001), and invasion range (*P* = 0.001), while it shows no correlation with gender, age, tumor size, or tumor differentiation (Table 2). These findings suggest that the m^6^A reader protein IGF2BP3 is highly expressed in GC and is closely associated with poor prognosis in GC patients.

**Figure 1.**
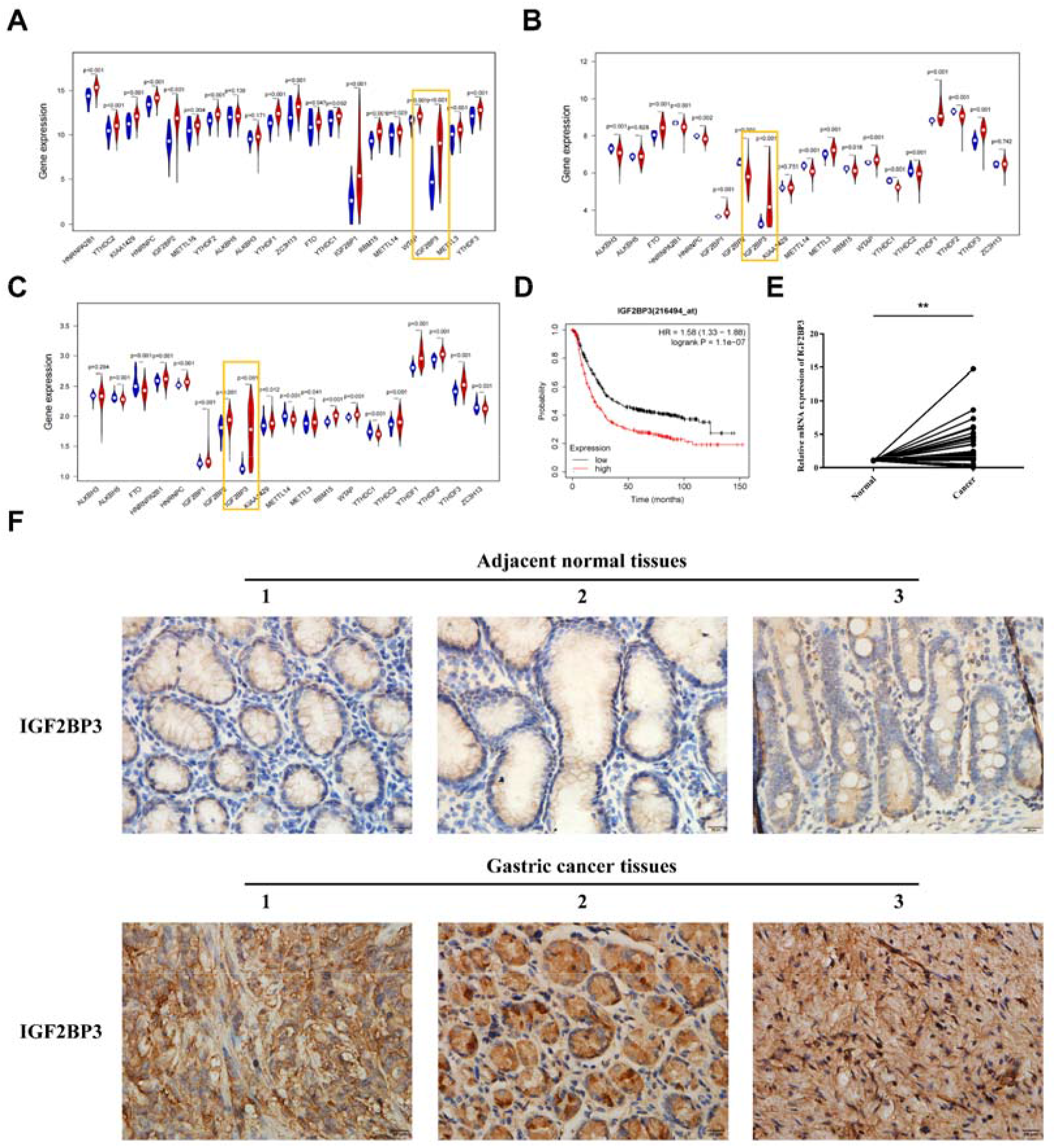
Increased IGF2BP3 expression is associated with poor prognosis in GC patients. **(A-C)** Expression of 21 m^6^A related genes in the TCGA database, as well as the GSE54129 and GSE66229 datasets. **(D)** Kaplan-Meier analysis of IGF2BP3 expression levels and overall survival in GC patients. **(E)** The expression levels of IGF2BP3 mRNA in 50 pairs of GC and adjacent normal tissue samples were detected by qRT-PCR. **(F)** Representative images showing IGF2BP3 protein expression levels in 63 pairs of GC and adjacent normal tissue samples, as detected by IHC. Scale bar, 20 μm. Data are presented as means ± SD. \*\**P* <0.01, \*\*\**P* <0.001

**Table 1.** The expression level of IGF2BP3 Protein in two groups.

| Group | Cases | IGF2BP3 expression | | Positive Rate(%) | $\chi^2$ | *P* |
| --- | --- | --- | --- | --- | --- | --- |
|  |  | High | Low |  |  |  |
| Gastric cancer tissues | 63 | 38 | 25 | 60% | 22.415 | < 0.001*** |
| Adjacent normal tissues | 63 | 12 | 51 | 19% |  |  |

**Table 2.** The relationship between the expression of IGF2BP3 in gastric cancer tissue and the clinicoapathological data of patients.

| Group | Cases | IGF2BP3 expression | | $\chi^2$ | *P* |
| --- | --- | --- | --- | --- | --- |
|  |  | High | Low |  |  |
| Gender |  |  |  |  |  |
| Male | 48 | 33 | 15 | 2.404 | 0.121 |
| Female | 15 | 7 | 8 |  |  |
| Age (years) |  |  |  |  |  |
| ≤60 | 16 | 11 | 5 | 0.256 | 0.613 |
| > 60 | 47 | 29 | 18 |  |  |
| Tumor size (cm) |  |  |  |  |  |
| < 5 | 34 | 20 | 14 | 1.550 | 0.213<sup>#</sup> |
| ≥ 5 | 27 | 20 | 7 |  |  |
| Differentiation |  |  |  |  |  |
| Well | 27 | 14 | 13 | 2.762 | 0.097 |
| Poorly | 36 | 26 | 10 |  |  |
| Lymphnode metastasis |  |  |  |  |  |
| Negative | 18 | 8 | 13 | 13.867 | < 0.001<sup>***</sup> |
| Positive | 45 | 35 | 10 |  |  |
| TNM Staging |  |  |  |  |  |
| 0 | 2 | 0 | 2 | 17.037 | 0.001<sup>**</sup> |
| I | 4 | 0 | 4 |  |  |
| II | 17 | 9 | 8 |  |  |
| III | 30 | 21 | 9 |  |  |
| IV | 10 | 10 | 0 |  |  |
| Invasion range |  |  |  |  |  |
| Tis | 2 | 0 | 2 | 14.165 | 0.001<sup>**</sup> |
| T1 | 3 | 0 | 3 |  |  |
| T2 | 8 | 5 | 3 |  |  |
| T3 | 5 | 1 | 4 |  |  |
| T4 | 45 | 34 | 11 |  |  |

### IGF2BP3 knockdown suppress the progression of GC *in vitro*

In order to explore the mechanism of IGF2BP3 in GC, we first downregulated IGF2BP3 expression in HGC-27 and MKN74 cell lines and observed the effects on various cellular functions. Following transfection with siRNA targeting IGF2BP3 (siIGF2BP3) and a siRNA negative control (siNC), both the mRNA (Figure 2A, B) and protein (Figure 2C, D) levels of IGF2BP3 were significantly reduced. The CCK-8 assay revealed that cells proliferation was markedly inhibited after knockdown of IGF2BP3 (Figure 2E, F). The wound healing assay (Figure 2G, H) and Transwell migration assay (Figure 2I, J) demonstrated that the downregulation of IGF2BP3 significantly suppressed cells migration. Furthermore, the Transwell invasion assay indicated that reducing IGF2BP3 expression significantly impaired the invasion capability of the cells (Figure 2K, L). Flow cytometry analysis showed that silencing IGF2BP3 expression promoted the apoptosis of HGC-27 and MKN74 cells (Figure 2M, N). These results suggest that downregulation of IGF2BP3 inhibits proliferation, migration and invasion of GC cells and promotes apoptosis.

**Figure 2.**
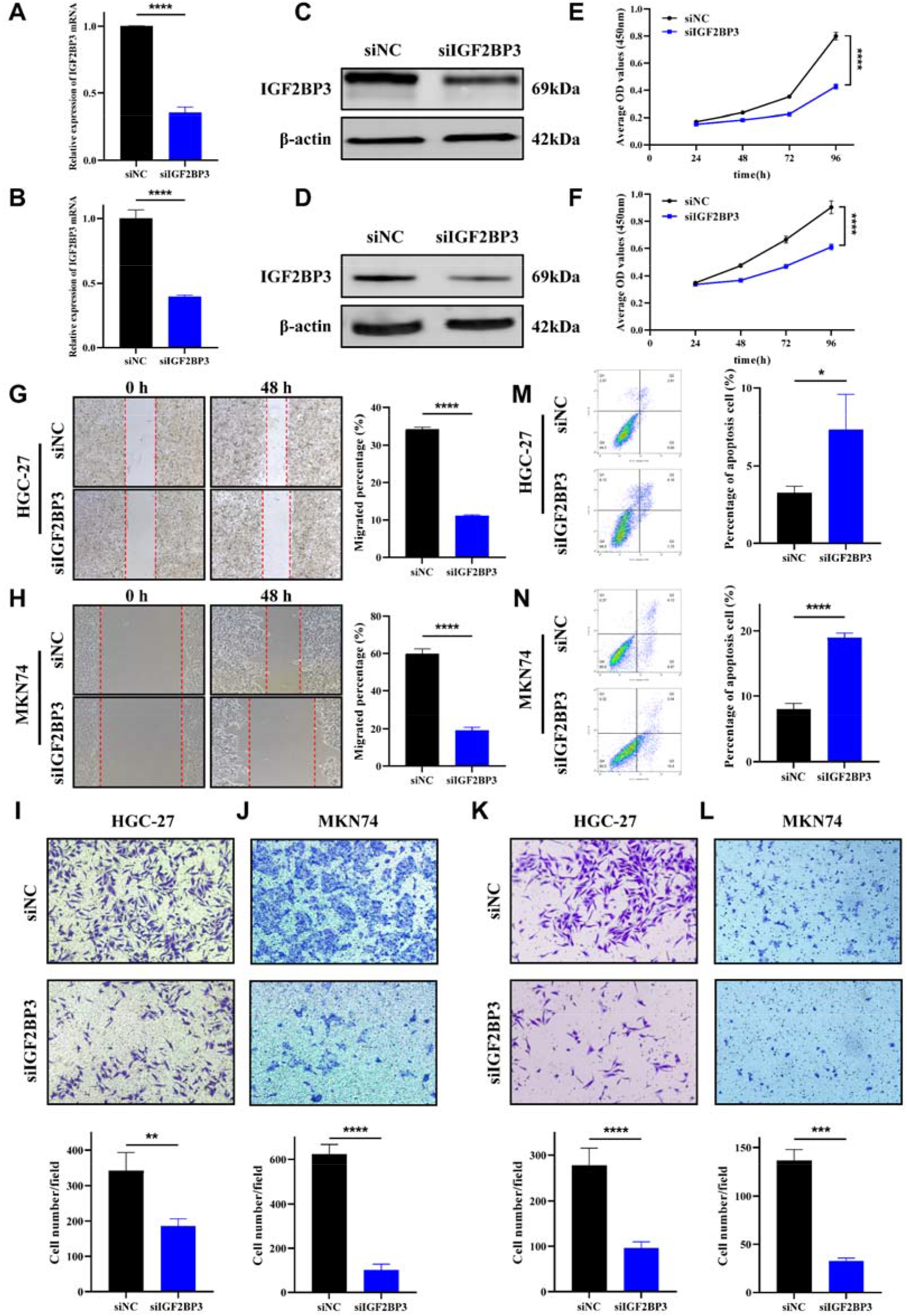
Knockdown of IGF2BP3 in GC cells suppresses cell proliferation, migration and invasion *in vitro*. **(A,B)** The expression of IGF2BP3 mRNA in HGC-27 and MKN74 cells was detected by qRT-PCR. **(C, D)** The expression of IGF2BP3 protein in HGC-27 and MKN74 was detected by WB. **(E, F)** The effect of IGF2BP3 knockdown on the proliferation of HGC-27 and MKN74 cells was determined by the CCK-8 assay. **(G, H)** The effect of IGF2BP3 knockdown on the migration of HGC-27 and MKN74 cells was assessed by the wound healing assay. Scale bar, 200 μm. **(I, J)** The effect of IGF2BP3 knockdown on the migration of HGC-27 and MKN74 cells was evaluated using the Transwell migration assay. Scale bar, 100 μm. **(K, L)** The effect of IGF2BP3 knockdown on the invasion of HGC-27 and MKN74 cells was assessed using Transwell invasion assay. Scale bar, 100 μm. **(M, N)** The effect of IGF2BP3 knockdown on the apoptosis of HGC-27 and MKN74 cells was determined by flow cytometry. Data are presented as means ± SD. \**P* <0.05, \*\**P* <0.01, \*\*\**P* <0.001, \*\*\*\**P* <0.0001

### Overexpression of IGF2BP3 can promote the progression of GC *in vitro*

We upregulated the expression of IGF2BP3 in MKN7 and GES-1 cell lines to observe its effect on cellular biological functions. After transfecting the cells with pcDNA3.1-IGF2BP3 (IGF2BP3) and pcDNA3.1-vector (Vector), we obversed a significant increase in IGF2BP3 mRNA (Figure 3A, B) and protein (Figure 3C, D) levels. The CCK-8 assay demonstrated a notable enhancement in cell proliferation following after the overexpression of IGF2BP3 (Figure 3E, F). Both the wound healing assay (Figure 3G, H) and the Transwell migration assay (Figure 3I, J) indicated that upregulating IGF2BP3 significantly strengthened the cells’ migration capability. Furthermore, the Transwell invasion assay revealed that elevated IGF2BP3 expression significantly heightened the invasion ability of cells (Figure 3K). The flow cytometry analysis revealed that increased IGF2BP3 expression led to a reduction in apoptosis levels in both MKN7 and GES-1 cell lines (Figure 3L, M). Despite multiple experiments, GES-1 cells did not demonstrate the ability to penetrate the matrix glue at the base of the Transwell chambers, due to the inherently weaker invasion ability of normal epithelial cells. In summary, these results show that overexpression of IGF2BP3 can promote the proliferation, migration and invasion of GC cells and inhibit their apoptosis, suggesting that IGF2BP3 may play the role of a pro-oncogene in GC.

**Figure 3.**
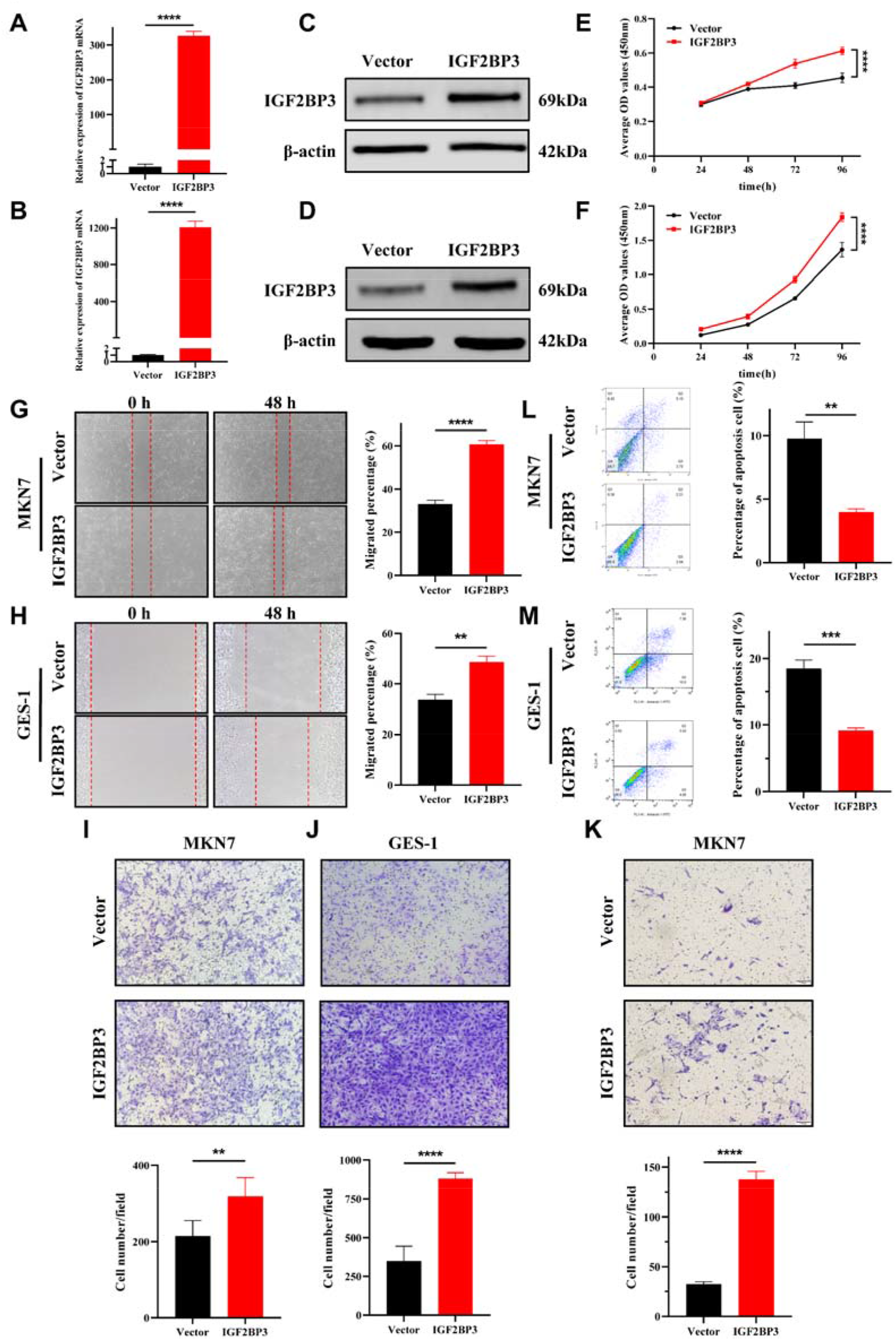
Overexpression of IGF2BP3 in GC cells and human gastric mucosa cells promotes cell proliferation, migration and invasion *in vitro*. **(A,B)** The expression of IGF2BP3 mRNA in MKN7 and GES-1 was detected by qRT-PCR. **(C, D)** The expression of IGF2BP3 protein in MKN7 and GES-1 was detected by WB. **(E, F)** The effect of IGF2BP3 overexpression on the proliferation of MKN7 and GES-1 cells was assessed using the CCK-8 assay. **(G, H)** The effect of IGF2BP3 overexpression on the migration of MKN7 and GES-1 cells was evaluated through a wound healing assay. Scale bar, 200 μm. **(I, J)** The effect of IGF2BP3 overexpression on the migration of MKN7 and GES-1 cells was determined utilizing a Transwell migration assay. Scale bar, 100 μm. **(K)** The effect of IGF2BP3 overexpression on the invasion of MKN7 cells was assessed via a Transwell invasion assay. Scale bar, 100 μm. **(L, M)** The effect of IGF2BP3 overexpression on the apoptosis of MKN7 and GES-1 cells was analyzed by flow cytometry. Data are presented as means ± SD. \*\**P* <0.01, \*\*\**P* <0.001, \*\*\*\**P* <0.0001

### IGF2BP3 promotes glucose metabolism in GC cells

Tumor cells rapidly produce a large amounts of energy through a unique aerobic glycolysis pathway to meet their surging metabolic demands, thereby maintaining normal cellular functions. We monitored the levels of glucose, lactic acid and ATP in GC cells after modulating the expression of IGF2BP3 to explore its role in glucose metabolism in these cells. The results showed that after down-regulating or up-regulating the expression level of IGF2BP3 in HGC-27 and MKN7 cells, the intracellular levels of glucose (Figure S1A), lactic acid (Figure S1B) and ATP (Figure S1C) were correspondingly decreased or increased. These findings suggest that IGF2BP3 may play a significant role in regulating glucose metabolism in GC cells. In the near future, we will continue to examine changes in the oxygen consumption rate (OCR) and extracellular acidification rate (ECAR) in GC cells after altering IGF2BP3 expression, to further explore the effects of IGF2BP3 on the glycolytic capability of these cells.

### IGF2BP3 deficiency inhibits the growth of GC *in vivo*

In order to further investigate the mechanism of IGF2BP3 in GC *in vivo*, we established tumor bearing nude mice models with low expression of IGF2BP3 and corresponding negative controls. HGC-27 cell lines stably transfected with shRNA-IGF2BP3 and shRNA-negative control were successfully established (shNC/shIGF2BP3) and subsequently injected subcutaneously into two groups of nude mice, respectively. The tumor-bearing models, along with the curves of tumor volume, tumor photographs and weights were presented in Figure 4A-D. The results demonstrated a significant reduction in the growth rate, volume, and weight of tumors in the shIGF2BP3 group compared to those in the shNC group. HE staining was utilized to evaluate the morphological characteristics of the xenograft tissues, and the results indicated that the cell morphology and structure were similar to those of GC tissues (Figure 4E). Ki67 immunohistochemical staining indicated that the proliferation ability of cells in the shIGF2BP3 group was significantly inhibited compared with the shNC group (Figure 4F). These findings suggest that IGF2BP3 knockdown can inhibit the growth of GC *in vivo*.

**Figure 4.**
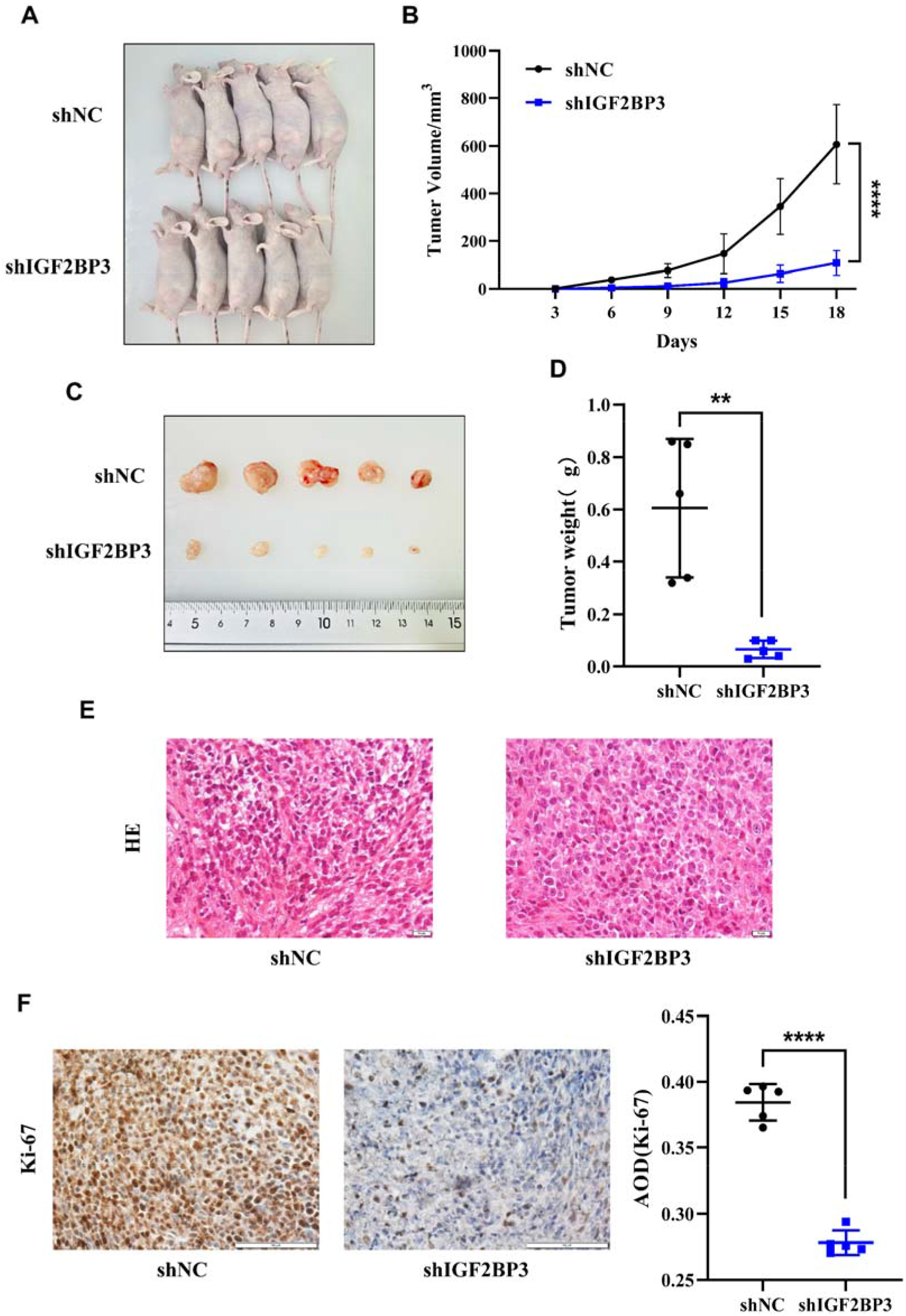
IGF2BP3 regulates the growth of GC *in vivo*. **(A)** Nude mouse tumor-bearing model. **(B)** Tumor growth curve. **(C, D)** Images (C) and weight (D) of xenografted tumors were analyzed. **(E)** Representative images of tumor tissue sections stained with HE. Scale bar, 50 μm. **(F)** Representative image of Ki67 IHC staining in tumor tissue sections. Scale bar, 50 μm. Data are shown as means ± SD. \*\**P* <0.01, \*\*\*\**P* <0.0001

### FBXO32 is the m^6^A modification target of IGF2BP3 in GC

In order to further explore the potential mechanism of IGF2BP3 in the pathogenesis of GC, we performed RNA-seq, MeRIP-seq, and RIP-seq in HGC-27 cells of siNC group and siIGF2BP3 group, respectively. The RNA-seq results indicated that the expression levels of a total of 388 genes were significantly changed after IGF2BP3 expression was knocked down, including 181 down-regulated genes and 207 up-regulated genes (Figure 5A, B). MeRIP-seq results showed that 25,331 m^6^A peaks were identified in the siNC group and 26,188 m^6^A peaks in the siIGF2BP3 group (Figure 5C). When mapping the m^6^A methyl group in HGC-27 cells, the m^6^A consistent sequence motif (GGAC) could be identified, indicating successful enrichment of mRNA with m^6^A modifications (Figure 5D). In addition, m^6^A modification was observed to be mainly enriched in the CDS and 3’ UTR of mRNA (Figure 5E). Then, the genes corresponding to the m^6^A peaks of the siNC group and siIGF2BP3 group were intersected, resulting in the identification of 10,514 genes (Figure 5F). Analysis of the RIP-seq results showed that m^6^A modification was also primarily enriched in the CDS and 3’ UTR of mRNA (Figure 5G). Specifically, 1,928 and 810 m^6^A peaks were identified in siNC group and siIGF2BP3 group, respectively (Figure 5H). After removing the same genes from the m^6^A peaks corresponding genes of both group, 73 unique genes were identified (Figure 5I). Subsequent unified analysis of the genes identified by RNA-seq, MeRIP-seq and RIP-seq showed that 19 genes bound to the IGF2BP3 protein were simultaneously labeled by m^6^A, and none of them had changes in transcription level after IGF2BP3 knockdown. These genes were identified as IGF2BP3 candidate target genes (Figure 5J). Further analysis of these genes included examing their expression in GC tissues, subcellular localization, survival of GC patients associated with these genes, and the abundance and loci of m^6^A on their mRNA. The results suggested that FBXO32 was highly expressed in GC tissues according to the TCGA database (Figure 5K), predominantly distributed in the cytoplasm and nucleus (Figure 5L), and the overall survival of patients with high expression of FBXO32 was poor (Figure 5M), and the IGF2BP3 binding site was coincide with the m^6^A modification site (Figure 5N). Additionally, RIP-qPCR demonstrated that the IGF2BP3 protein could significantly enrich FBXO32 mRNA (Figure 5O). Moreover, as predicted by the catRAPID website, there is a direct binding site between the IGF2BP3 protein and FBXO32 mRNA, as shown in the motif (Figure 5P). Based on the above analysis results, FBXO32 was identified as the m^6^A modification target of IGF2BP3 in GC.

**Figure 5.**
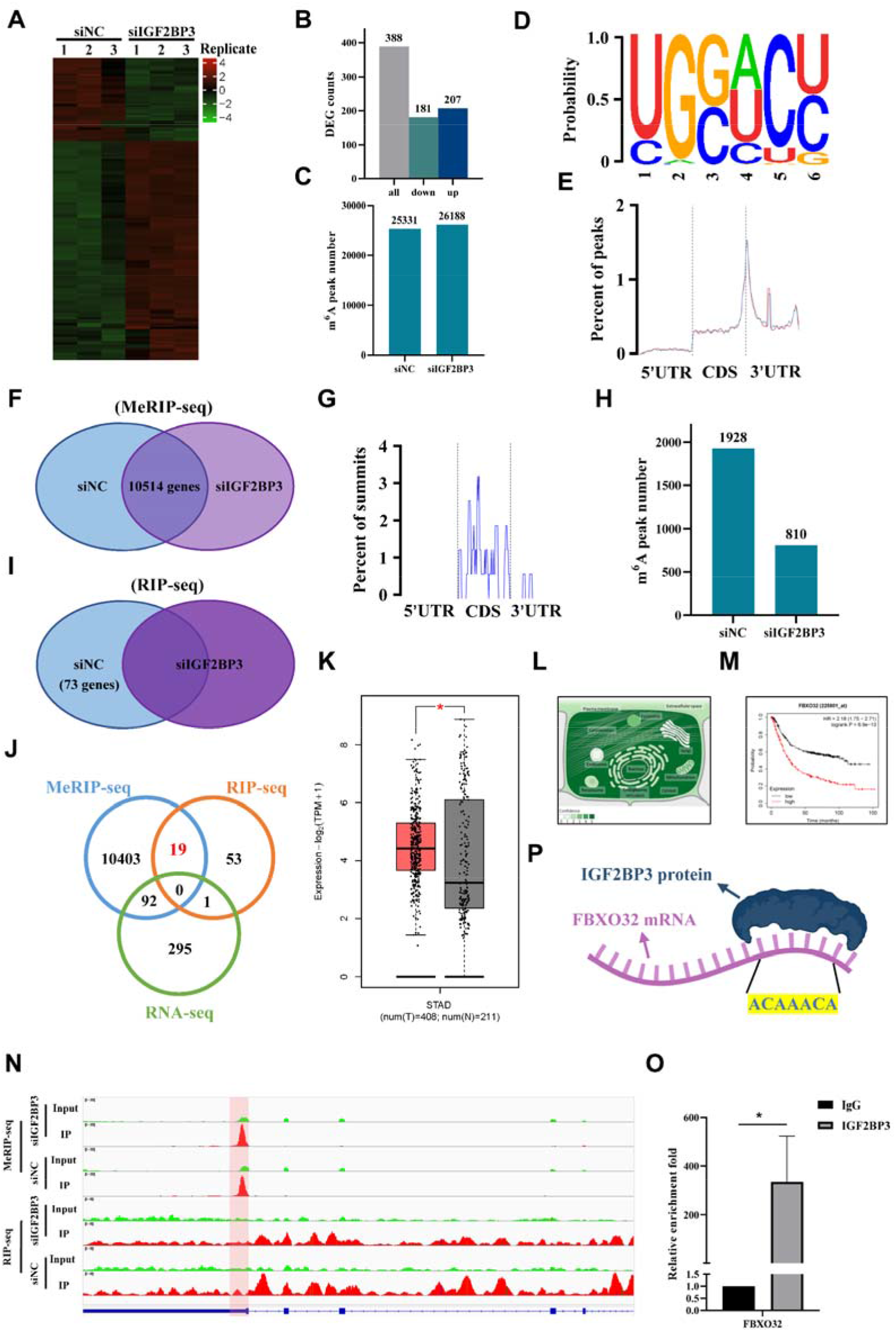
FBXO32 is the m^6^A modification target of IGF2BP3. **(A)** Differential expression genes (DEGs) heat map identified by RNA-seq. **(B)** The number of DEGs identified by RNA-seq. **(C)** The number of m^6^A peaks identified by MeRIP-seq. **(D)** m^6^A motif identified by MeRIP-seq. **(E)** The enrichment regions of m^6^A peaks on mRNA identified by MeRIP-seq. **(F)** The common genes between the siNC group and the siIGF2BP3 group in MeRIP-seq. **(G)** The enrichment regions of m^6^A peaks on mRNA identified by RIP-seq. **(H)** The number of m^6^A peaks identified by RIP-seq. **(I)** Genes identified by RIP-seq. **(J)** Overlap analysis of MeRIP-seq, RIP-seq and RNA-seq identified genes. **(K)** Expression of FBXO32 in GC paired tissue cohort from the TCGA database. **(L)** Subcellular localization of FBXO32 from GeneCards. **(M)** Kaplan-Meier analysis of FBXO32 expression levels and overall survival in patients with GC. **(N)** The m^6^A site and its abundance on FBXO32 transcripts were analyzed using IGV software. **(O)** The interaction between IGF2BP3 and FBXO32 was detected by RIP-qPCR. **(P)** Schematic diagram of the catRAPID website predicting the binding site between IGF2BP3 protein and FBXO32 mRNA.

### FBXO32 is highly expressed in GC and directly interacts with IGF2BP3

Next, we initially verified the expression of FBXO32 in GC cell lines and GC tissues. The results indicated that the level of FBXO32 mRNA in GC cell lines was higher than that in normal gastric mucosa cells (Figure 6A), and in GC tissues, it was higher in adjacent tissues (34 pairs of GC paired tissues) (Figure 6B). The expression of FBXO32 protein was highest in MKN7 cells (Figure 6C), and higher in GC tissues than in adjacent tissues (19 pairs of GC paired tissues) (Figure 6D). These results suggest that FBXO32 may acts as an oncogene in GC pathogenesis. Subsequently, the interaction between IGF2BP3 and FBXO32 was verified by Co-IF experiments. The Co-IF results showed that IGF2BP3 and FBXO32 were mainly located in the cytoplasm and nucleus of GC cells, and they have co-localization (Figure 6E). Finally, to further confirm the regulation of FBXO32 expression by IGF2BP3 in GC cells, we examined the transcriptional and translational changes of FBXO32 after IGF2BP3 knockdown or overexpression. After down-regulating the expression level of IGF2BP3 in HGC-27 cells, the expression level of FBXO32 mRNA did not change significantly (Figure 6F), while the expression level of its protein was down-regulated (Figure 6G). Similarly, when IGF2BP3 expression was upregulated in MKN7 cells, the expression level of FBXO32 mRNA was not significantly changed (Figure 6H), while its protein expression level was upregulated (Figure 6I). Theses phenomena indicate that IGF2BP3, as an m^6^A reader, only regulates the expression of downstream genes at the translation level, not at the transcriptional level, which is consistent with the existing research conclusions on other m^6^A readers. In conclusion, these results indicate that IGF2BP3 and FBXO32 interact with each other in GC, and IGF2BP3 can regulate the expression of FBXO32 protein.

**Figure 6.**
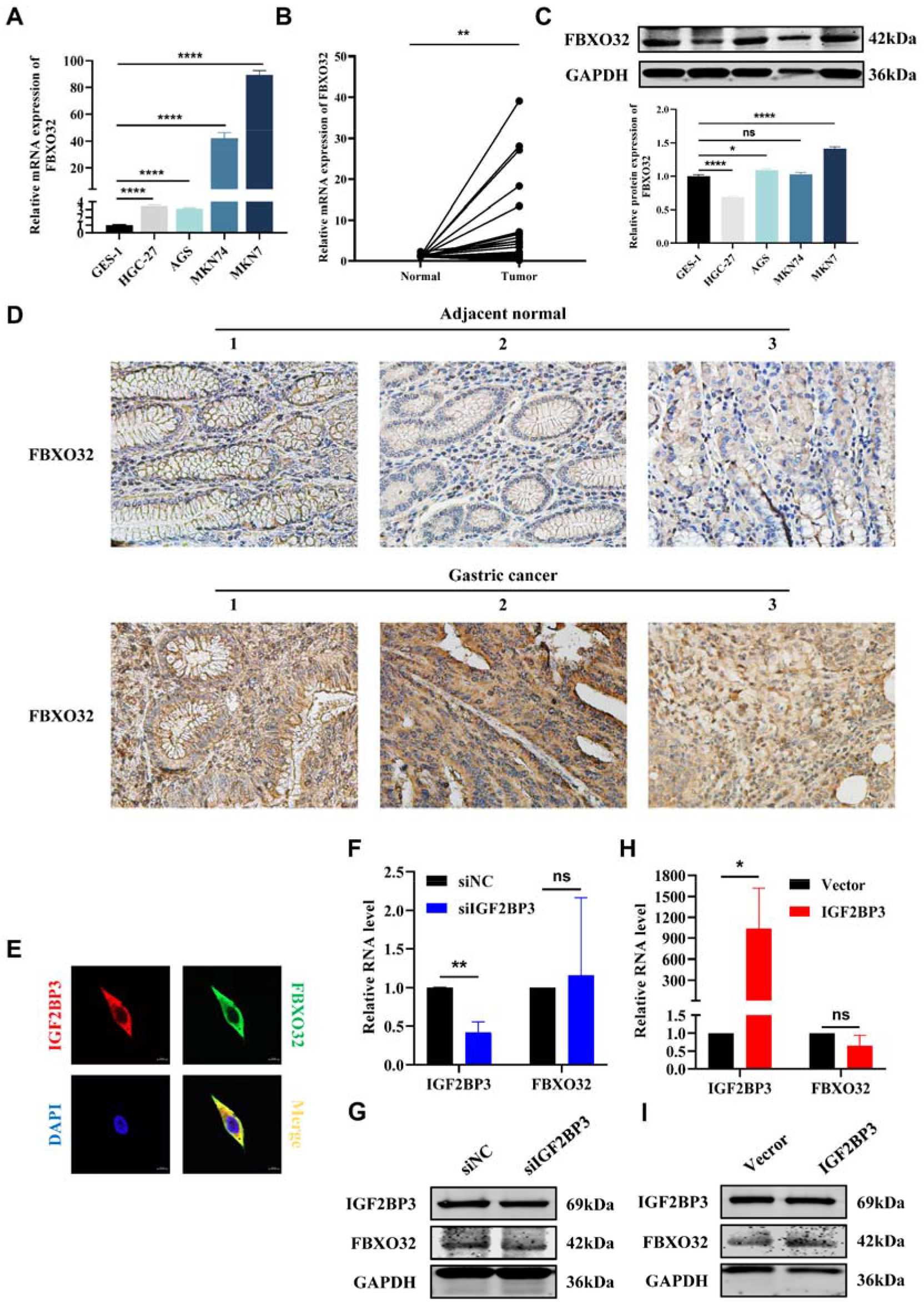
Expression of FBXO32 and its interaction with IGF2BP3 in GC. **(A)** The expression of FBXO32 mRNA in GC cell lines and GES-1 was detected by qRT-PCR. **(B)** The expression of FBXO32 mRNA in 34 pairs of GC paired tissues was detected by qRT-PCR. **(C)** The basal expression of FBXO32 protein in cell lines was detected by WB. **(D)** The expression of FBXO32 protein in 19 pairs of GC paired tissues was detected by IHC. Scale bar, 50 μm. **(E)** The distribution of IGF2BP3 and FBXO32 in HGC-27 cells was confirmed by Co-IF. Scale bar, 10 μm. **(F)** qRT-PCR was used to detect FBXO32 mRNA expression after IGF2BP3 knockdown in HGC-27 cells. **(G)** WB was used to detect FBXO32 protein expression after IGF2BP3 knockdown in HGC-27 cells. **(H)** qRT-PCR was used to detect FBXO32 mRNA expression after IGF2BP3 overexpression in MKN7 cells. **(I)** WB was used to detect FBXO32 protein expression after IGF2BP3 overexpression in MKN7 cells. Data are presented as means ± SD. \**P* <0.05, \*\**P* <0.01, \*\*\*\**P* <0.0001

### The oncogenic effect of IGF2BP3 depends on targeting FBXO32 and activating the cGMP-PKG signaling pathway

Four siRNA-FBXO32 sequences were transfected simultaneously into MKN7 cells with the highest FBXO32 expression, and one of the siRNA sequences with the most significant knockdown efficiency (si-FBXO32-910) was screened for subsequent experiments (Figure S2A). Next, we transfected siRNA-negative control/siRNA-IGF2BP3 and pcDNA3.1-vector/pcDNA3.1-FBXO32 into HGC-27 and pcDNA3.1-vector/pcDNA3.1-IGF2BP3 and siRNA-negative control/siRNA-FBXO32 into MKN7 cells, respectively or simultaneously. The changes of IGF2BP3 expression in each group were detected by qRT-PCR. The results suggest that the knockdown effect of IGF2BP3 in HGC-27 cells could be attenuated by the overexpression of FBXO32 (Figure S2B), and the up-regulation effect of IGF2BP3 in MKN7 cells could also be weakened by the knockdown of FBXO32 (Figure S2C). These findings indicate that FBXO32 can affect the expression of IGF2BP3 in GC cells. To further explore the downstream molecular mechanisms of FBXO32, we selected the top 1000 genes co-expressed with FBXO32 in gastric adenocarcinoma from the TCGA database and analyzed the signaling pathways controlled by them. The results suggested that genes in the cGMP-PKG signaling pathway were significantly enriched (Figure 7A). Subsequently, we tested whether FBXO32 could rescue the inhibitory effects on cGMP-PKG signaling pathway activation induced by the knockdown of IGF2BP3. The results suggest that when the expression level of IGF2BP3 is downregulated in HGC-27 cells, the expression levels of downstream effectors of cGMP (PKG1, pVASP) are also significantly downregulated. Concurrently, overexpression of FBXO32 can rescue the downregulatory effects on these effectors caused by the knockdown of IGF2BP3 (Figure 7B). These findings suggest that FBXO32 is involved in the regulation of IGF2BP3’s activation of the cGMP-PKG signaling pathway. Next, we further explored whether the effects of IGF2BP3 on GC cell function would be influenced by FBXO32. The results showed that the attenuation of cell proliferation, migration, and invasion induced by knockdown of IGF2BP3 in HGC-27 cells could be rescued by the up-regulation of FBXO32 expression (Figure 7C-G). The enhancement of cell proliferation, migration, and invasion caused by overexpression of IGF2BP3 in MKN7 cells could be attenuated by the down-regulation of FBXO32 expression (Figure 7D-L). In summary, FBXO32 serves as a crucial downstream target for IGF2BP3 in promoting the progression of GC, and the oncogenic effect of IGF2BP3 in GC is dependent on FBXO32 and the cGMP-PKG signaling pathway. Therefore, we propose that increased IGF2BP3 can bind to more FBXO32 mRNA by recognizing m^6^A sites on the FBXO32 mRNA, leading to an increase in FBXO32 protein expression and subsequently activating the cGMP-PKG signaling pathway, ultimately promoting tumorigenesis in GC cells (Figure 7M).

**Figure 7.**
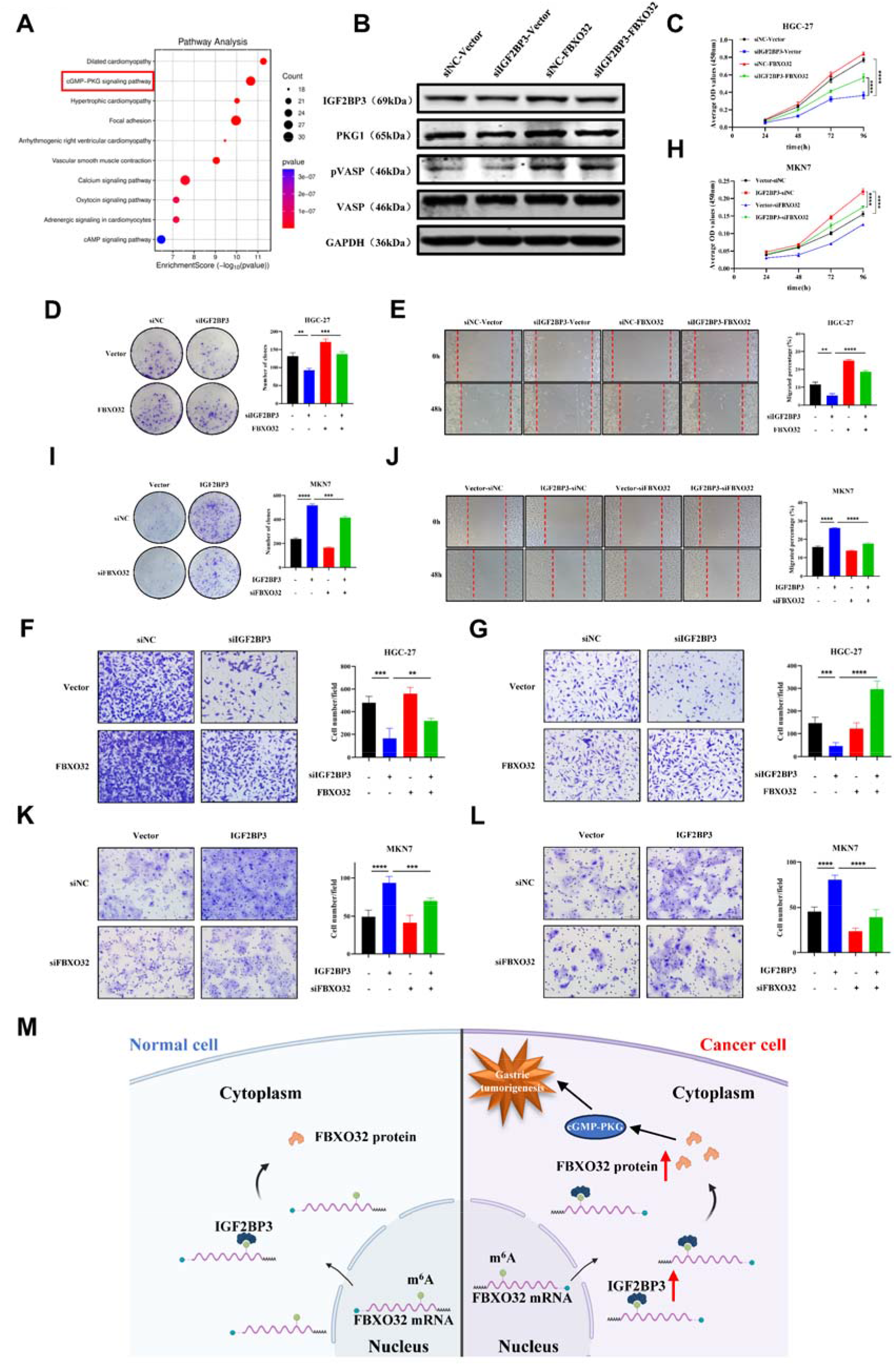
Oncogenic function of IGF2BP3 depends on FBXO32 and cGMP-PKG signaling pathway. **(A)** KEGG enrichment result of the top 1000 genes coexpressed with FBXO32 in gastric adenocarcinoma in TCGA database. **(B)** The expression levels of cGMP-PKG signaling pathway related proteins were detected by WB after instantaneous co-transfection of IGF2BP3 siRNA and FBXO32 overexpression plasmid in HGC-27. **(C-G)** The effects of FBXO32 on IGF2BP3 in HGC-27 cells were detected by CCK-8 assay (C), colony formation assay (D), wound healing assay (E), Scale bar, 200 μm, Transwell migration assay (F) and Transwell invasion assay (G). Scale bar, 100 μm. **(H-L)** The effects of FBXO32 on IGF2BP3 in MKN7 cells were detected by CCK-8 assay (H), colony formation assay (I), wound healing assay (J), Scale bar, 200 μm, Transwell migration assay (K) and Transwell invasion assay (L). Scale bar, 100 μm. **(M)** Schematic illustration of the molecular mechanism of IGF2BP3 in GC. Data are presented as means ± SD. \**P* <0.05, \*\**P* <0.01, \*\*\**P* <0.001, \*\*\*\**P* <0.0001

## Discussion

In this study, we report for the first time that IGF2BP3 binds to FBXO32 mRNA by recognizing the m^6^A site, resulting in increased intracellular level of FBXO32 protein, which then activates the cGMP-PKG signaling pathway, ultimately promotes the progression of GC. Currently, no optimal prognostic biomarker for GC has been identified. This study also confirmed that the expression of IGF2BP3 is upregulated in GC, which is associated with poor prognosis in patients and alterations in cellular glucose metabolism. It has unveiled the clinical significance and potential value of the IGF2BP3/FBXO32/cGMP-PKG axis in the therpy of GC.

Through literature review and database analysis, we found that the expression of IGF2BP3 was most significantly different in GC tissues compared to adjacent normal tissues. However, the relationship between IGF2BP3 and GC remains to be further explored. Previous studies indicate that the expression level of IGF2BP3 in laryngeal squamous cell carcinoma (LSCC) was higher than that in adjacent normal tissues^[15]^. Additionally, the expression of IGF2BP3 is elevated in gallbladder cancer and is associated with poor prognosis of patients, which can enhance multiple malignant phenotypes of gallbladder cancer cells, and knockdown of IGF2BP3 can inhibit the tumor growth *in vivo*^[19]^. The expression level of IGF2BP3 in colon cancer is also significantly higher than that in adjacent normal tissues and is associated with worse overall survival. Altering the expression level of IGF2BP3 can regulate the cell cycle and affect cell proliferation, and knocking it down can inhibit the tumor growth of colon cancer *in vivo*^[17]^. Consistent with these findings, our study revealed that the expression levels of IGF2BP3 mRNA and protein in GC tissues were higher than those in adjacent normal tissue. The increased expression level of IGF2BP3 was closely related to lymph node metastasis, more advanced TNM stage and deeper invasion depth. In our experiments, knocking down the expression level of IGF2BP3 in GC cells significantly inhibited, cell proliferation, migration and invasion, and increased apoptosis. Conversely, upregulating IGF2BP3 expression in GC cells had the opposite effect. Furthermore, we observed that IGF2BP3 deficiency inhibited the growth of GC *in vivo*. These findings suggest that IGF2BP3 may be an important oncogene in GC. Additionally, Wang J et al. found that IGF2BP2 enhanced the stability of GLUT1 mRNA, a key gene of aerobic glycolysis, in an m^6^A dependent manner, significantly promoting the aerobic glycolysis in colorectal cancer cells^[18]^. Our results also showed that after up-regulating or down-regulating the expression level of IGF2BP3 in GC cells, the contents of glucose, lactic acid and ATP in the cells correspondingly increased/decreased. Thus, the effects of IGF2BP3 on the biological function of GC cells may be closely related to the regulation of glucose metabolism in cells, which warrants further exploration.

Through RNA-seq, MeRIP-seq, RIP-seq and bioinformatics analysis, we identified FBXO32 as the target of IGF2BP3 with m^6^A modification in GC for the first time. F-Box Protein 32 (FBXO32), originally identified as a muscle-specific gene involved in muscle atrophy, is an E3 ubiquitin ligase located on chromosome 8q24.13^[20, 21]^. Previous studies have reported that FBXO32 is involved in various processes such as epithelial-mesenchymal transformation and tumorigenesis^[22]^. The overexpression of FBXO32 is conducive to the proliferation and migration of melanoma cells, while its downregulation has the opposite effect^[23]^. FBXO32 is abnormally upregulated in pancreatic ductal adenocarcinoma, promoting the migration and invasion of pancreatic cancer cells *in vitro*, and promoting tumor growth and metastasis *in vivo*^[24]^. FBXO32 enhances the PI3K/AKT/mTOR pathway by degrading PTEN, thereby promoting the progression of lung adenocarcinoma^[25]^. In this study, the expression levels of FBXO32 mRNA and protein in GC tissues were higher than those in adjacent normal tissues, and the expression levels of FBXO32 mRNA and protein in GC cells were also higher than those in normal gastric mucosa cells. These findings suggest that FBXO32 may be an important oncogene in GC. However, there are some results that contradict our conclusions. For example, the expression of FBXO32 is down-regulated in multiple myeloma^[26]^. Overexpression of FBXO32 can inhibit tumorigenicity of ovarian cancer cells both *in vivo* and *in vitro*^[27]^. Therefore, the role of FBXO32 in cancer cannot be generalized, and more extensive and in-depth studies are needed to elucidate its mechanisms. Subsequently, Co-IF experiments confirmed that IGF2BP3 and FBXO32 had a direct interaction relationship in GC cells, and this conclusion has not been reported in existing studies. In addition, as predicted by bioinformatics analysis, there was a binding site between IGF2BP3 protein and FBXO32 mRNA, which further supports our conclusion about their interaction. To explore the regulatory mechanism of IGF2BP3 on FBXO32 in GC, we demonstrated that IGF2BP3 regulates FBXO32 expression at the translational level but not at the transcriptional level. Previous studies have reported that IGF2BP3 can enhance the translation of COPS7B mRNA and promote the growth and metastasis of colorectal cancer^[28]^. IGF2BP3 promotes EGFR mRNA translation in an m^6^A-dependent manner, contributing to CRC progression and cetuximab resistance^[29]^. Thus, it is possible that IGF2BP3 binds FBXO32 mRNA in an m^6^A-dependent manner and promotes its translation, ultimately contributing to GC progression.

To further elucidate the mechanism by which IGF2BP3 targets FBXO32 to promote GC progression, we analyzed the signaling pathways controlled by the top 1000 genes co-expressed with FBXO32 in gastric adenocarcinoma from the TCGA database. The results suggest that these genes are significantly enriched in the cGMP-PKG signaling pathway. The cGMP-PKG signaling pathway was originally reported to participate in the inhibition of platelet activation and the phosphorylation of VASP regulated by this pathway is an integral part of an efficient and sensitive signaling cascade^[30]^. Many recent studies have shown that the cGMP-PKG signaling pathway is also involved in the development of various cancers. For example, cGMP can activate PKG and downstream MAPK pathways, resulting in the increased tumor cell stemness and metastasis^[31]^; DARS-AS1 can accelerate the progression of cervical cancer by activating the cGMP-PKG pathway^[32]^; NO-cGMP-PKG can increase the proliferation of colon cancer cells while inhibiting apoptosis by activating ERK-1/2 and AP-1^[33]^. Subsequently, we confirmed through WB that the inhibitory effect on cGMP-PKG signaling pathway activation caused by the knockdown of IGF2BP3 can be rescued by FBXO32, and FBXO32 can also activate the cGMP-PKG signaling pathway. Finally, we also conducted rescue studys to further confirm that the IGF2BP3/FBXO32/cGMP-PKG axis can affect the proliferation, migration, and invasion capabilities of GC cells. These results indicate that the IGF2BP3/FBXO32/cGMP-PKG axis is involved in the occurrence and progression of GC and represents a promising new therapeutic target for clinical treatment of GC.

Admittedly, our research currently has certain limitations. For example, the human tissue samples used for clinical validation were all from the same hospital, and there was a lack of multi-center clinical data. In addition, our findings about the effect of changes in IGF2BP3 expression level on cellular glucose metabolism in GC cells are worthy of further exploration to elucidate the in-depth mechanism of GC and glucose metabolism.

In conclusion, our study demonstrated that IGF2BP3 is a oncogenic protein in GC, which promotes the translation of FBXO32 in an m^6^A dependent manner and activates the downstream cGMP-PKG signaling pathway, thereby modulating multiple biological functions of GC cells and ultimately promoting the progression of GC. Consequently, targeting the IGF2BP3/FBXO32/cGMP-PKG axis may offer a novel approach to the clinical treatment of GC.

## Acknowledgments

This work was supported by Hebei Provincial Government-funded Provincial Medical Excellent Talent Project (ZF2023025, ZF2024134, and LS202008), Hebei Natural Science Foundation (H2022206292, H2024206140), Key R&D Program of Hebei Province (223777103D and 223777113D), Prevention and treatment of geriatric diseases by Hebei Provincial Department of Finance (LNB202202, LNB201809 and LNB201909), Spark Scientific Research Project of the First Hospital of Hebei Medical University (XH202312 and XH201805), Hebei Province Medical Applicable Technology Tracking Project (G2019035) and other projects of Hebei Province (1387 and SGH201501).

## CRediT authorship contribution statement

**Yi Si:** Conceptualization, Data curation, Formal analysis, Methodology, Validation, Visualization, Writing – original draft, Writing – review & editing. **Bo Tian:** Conceptualization, Data curation, Formal analysis, Visualization, Writing – original draft, Writing – review & editing. **Rui Zhang:** Visualization, Writing – original draft. **Mingda Xuan:** Methodology, Writing – original draft. **Kunyi Liu:** Visualization, Writing – original draft. **Jiao Jiao**: Data curation, Writing – original draft. **Shuangshuang Han:** Formal analysis, Visualization, Writing – original draft. **Hongfei Li:** Formal analysis, Writing – original draft. **Yanhong Hu:** Formal analysis, Writing – original draft. **Hongyan Zhao:** Investigation, Writing – original draft. **Wenjing He:** Investigation, Writing – original draft. **Jia Wang:** Conceptualization, Funding acquisition, Project administration, Resources, Supervision, Writing – review & editing. **Ting Liu:** Conceptualization, Funding acquisition, Project administration, Validation, Writing – review & editing. **Weifang Yu:** Conceptualization, Funding acquisition, Project administration, Resources, Supervision, Writing – review & editing.

## Conflict Of Interest Statement

The authors have no conflicts of interest to declare.

## Supplementary Table

Supplementary Table 1 The primer sequences of IGF2BP3, FBXO32, β-actin.

## Supplementary Figures

**Supplementary Figure 1. IGF2BP3 affects glucose metabolism in GC cell lines. (A)** The effect of IGF2BP3 expression level on glucose uptake in GC cells was determined using a glucose uptake assay. **(B)** The effect of IGF2BP3 expression level on lactic acid production in GC cells was determined by utilizing a lactic acid production assay. **(C)** An ATP content assay showed the effect of IGF2BP3 expression level on ATP content in GC cells.

**Supplementary Figure 2. The knockdown efficiency of FBXO32 and the expression level of IGF2BP3 were detected by qRT-PCR. (A)** The knockdown efficiency of four siRNA-FBXO32 sequences in MKN7 cells was assessed by qRT-PCR. **(B, C)** The expression of IGF2BP3 mRNA in HGC-27 (B) and MKN7 (C) cells was measured by qRT-PCR after co-transfection of respective siRNAs and overexpression plasmids.

**Supplementary Figure 3. Protein interaction network diagram.**

## Notes

### Competing Interest Statement

The authors have declared no competing interest.

### Summary of Updates

Figure 6 revised Figure 7 revised Supplemental files updated

