## Supplemental Data 1 for "IGF2BP3 promotes the progression of gastric cancer by activating cGMP-PKG signaling pathway via targeting FBXO32"

**Supplementary Table 1 The sequence of primers**

| Gene | Forward Primer Sequence 5’-3’ | Reverse Primer Sequence 5’-3’ |
|---|---|---|
| IGF2BP3 | ACTGCACGGGAAACCCATAG | CCAGCACCTCCCACTGTAAAT |
| FBXO32 | TACGTGGTCCGGCTGTTG | CCATCCGATACACCCACATG |
| β-actin | CTCCATCCTGGCCTCGCTGT | GCTGTCACCTTCACCGTTCC |

**Supplementary Figure 1**

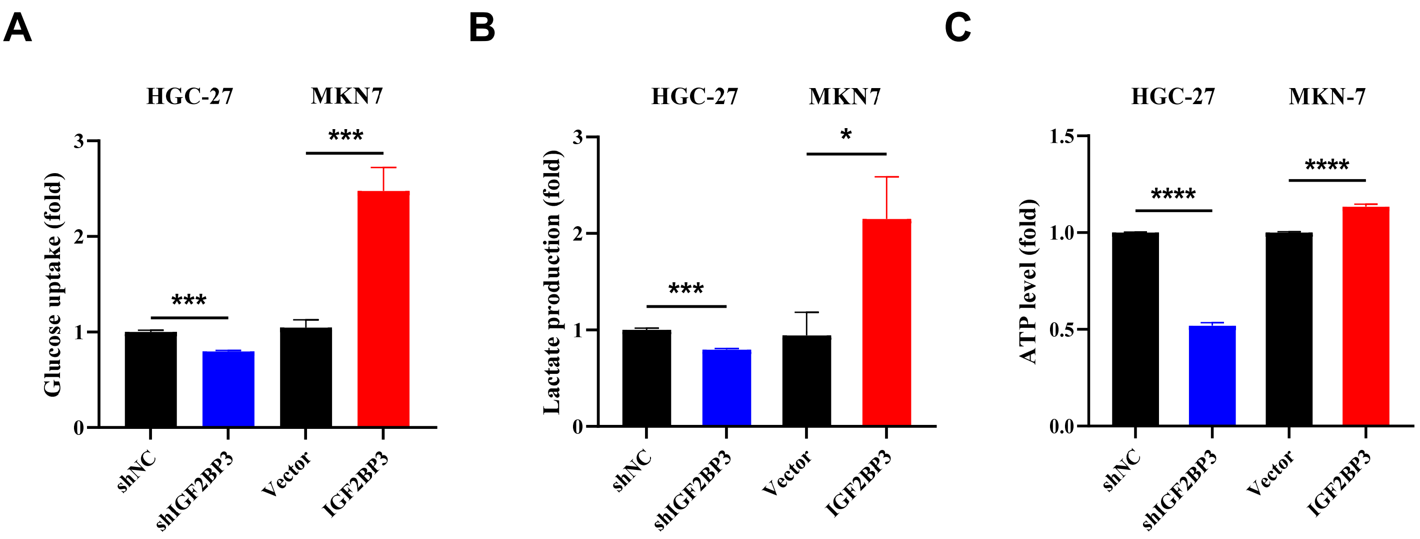


**Supplementary Figure 2**

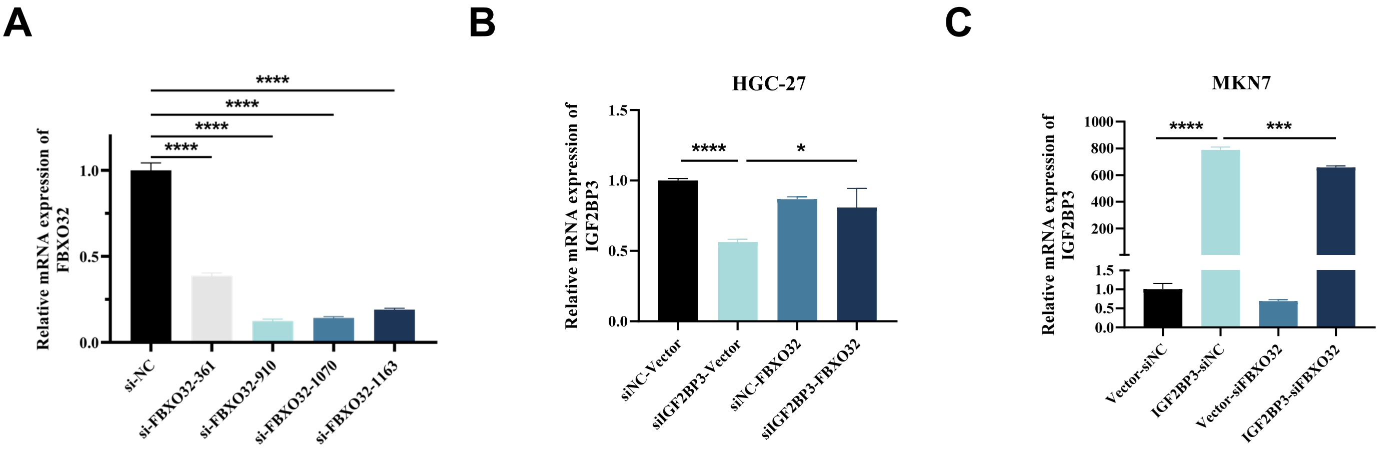


**Supplementary Figure 3**

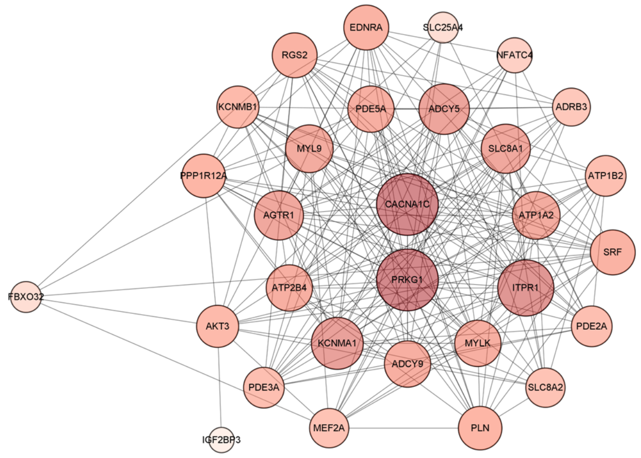
